# SpheroidAnalyseR – an online platform for analysing data from 3D spheroids or organoids grown in 96-well plates

**DOI:** 10.1101/2022.02.18.481039

**Authors:** Rhiannon Barrow, Joseph N Wilkinson, Yichen He, Martin Callaghan, Anke Brüning-Richardson, Mark Dunning, Lucy F Stead

## Abstract

Spheroids and organoids are increasingly popular three-dimensional (3D) cell culture models. Spheroid models are more physiologically relevant to a tumour compared to two-dimensional (2D) cultures and organoids are a simplified version of an organ with similar composition. Both can be used in cell migration studies, disease modelling and drug discovery. A drawback of these models is, however, the lack of appropriate analytical tools for high throughput imaging and analysis over a time course. To address this, we have developed an R Shiny app called SpheroidAnalyseR: a simple, fast, effective open-source app that allows the analysis of spheroid or organoid size data generated in a 96-well format. SpheroidAnalyseR processes and analyses datasets of image measurements that can be obtained via a bespoke software, described herein, that automates spheroid imaging and quantification using the Nikon A1R Confocal Laser Scanning Microscope. However, templates are provided to enable users to input spheroid image measurements obtained by user-preferred methods. SpheroidAnalyseR facilitates outlier identification and removal followed by graphical visualisation of spheroid measurements across multiple predefined parameters such as time, cell-type and treatment(s). Spheroid imaging and analysis can, thus, be reduced from hours to minutes, removing the requirement for substantial manual data manipulation in a spreadsheet application. The combination of spheroid generation in 96-well ultra-low attachment microplates, imaging using our bespoke software, and analysis using SpheroidAnalyseR toolkit allows high throughput, longitudinal quantification of 3D spheroid growth whilst minimising user input and significantly improving the efficiency and reproducibility of data analysis. Our bespoke imaging software is available from https://github.com/GliomaGenomics. SpheroidAnalyseR is available at https://spheroidanalyser7.azurewebsites.net, and the source code found at https://github.com/GliomaGenomics.

Graphical abstract - SpheroidAnalyseR main steps.
A summary of the process followed within the SpheroidAnalyseR app. Dotted boxes are the steps required to produce the input data for SpheroidAnalyseR. Solid boxes show the 5 main steps of SpheroidAnalyseR: Data input, Preview, Outlier removal, Merging and Plotting.

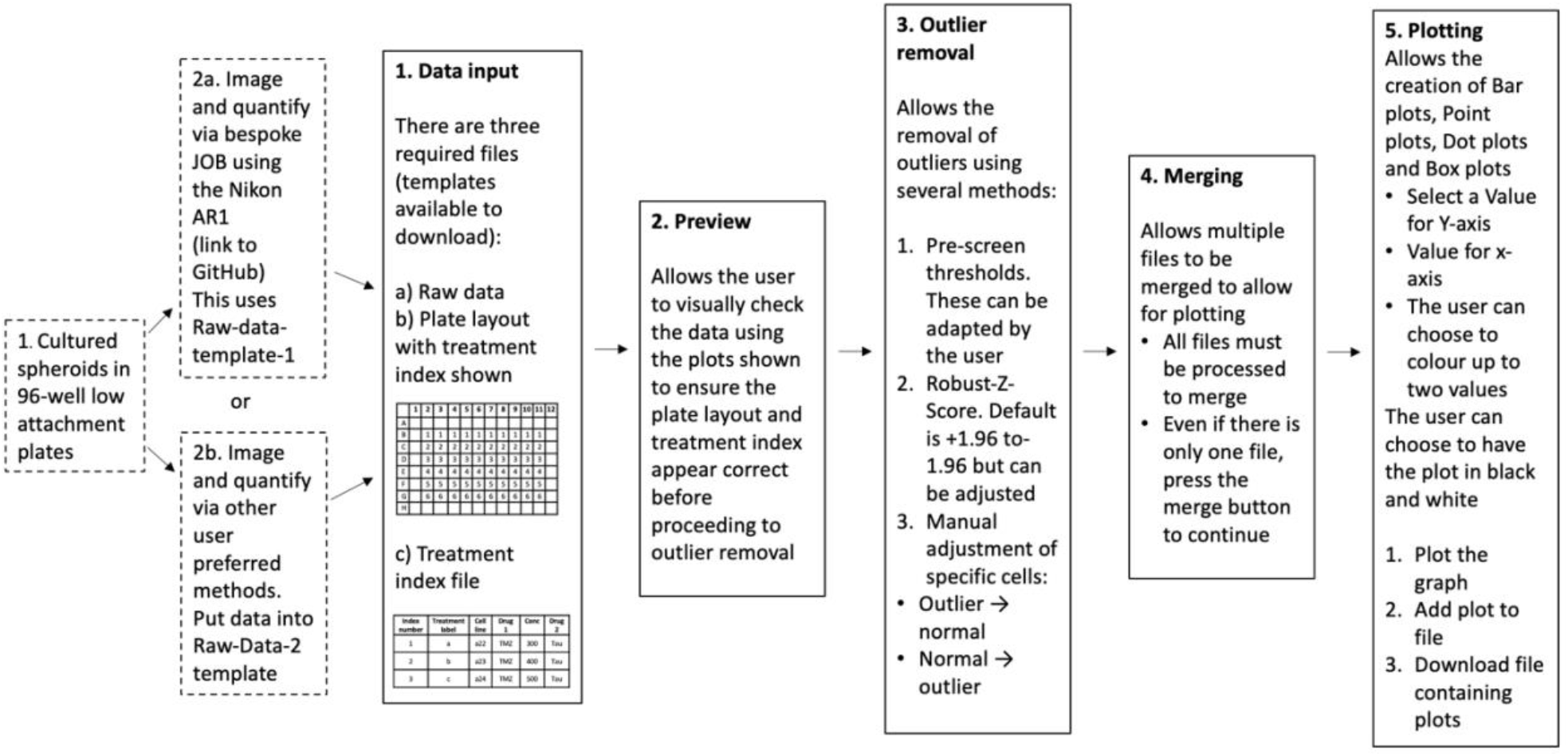

## BACKGROUND

### Spheroids and organoids as 3D cell culture models

The use of three-dimensional (3D) models such as spheroid and organoids has become an intermediate step between two-dimensional 2D cell culture and *in vivo* models especially in cancer research. 2D cell culture models have been used since the 19^th^ Century and still play a important role in scientific research due to their ease of use and readily available assays [1]. However, often the results from 2D models do not reflect cellular activity and responses as observed *in vivo*. Spheroid models are based on the generation of 3D structures of tumour cells and provide a more physiologically relevant environment for cancer cells in comparison to being grown as 2D monolayers; as such they lead to a more clinically relevant recapitulation of the biology or therapeutic response observed *in vivo* [2] and thus make a case for use as enhanced models in numerous drug discovery and toxicity studies [3]. As miniature versions of organs, organoids, typically grown from stem cells, are often used in disease modelling, drug discovery and cell migration studies [4] [5]. Cells contained within a 3D structure have many more physiological relevant features than cells grown as monolayers. For example, in 3D cells display a more natural morphology rather than being elongated, enabling them to grow in aggregates as opposed to a monolayer and they are exposed to varying oxygen and nutrient gradients. This allows them to be in different stages of the cell cycle and have more biologically relevant cell-cell interactions [6] [7]. The presence of an oxygen gradient means that tumour spheroids often have a hypoxic core which is a common feature in most solid tumour types such as glioblastoma (GBM) and breast cancer resembling the features of a tissue or tumour in a way that 2D cell cultures cannot. Furthermore, spheroids give a more accurate representation of drug response, and gene and protein expression profiles often show increased similarity to *in vivo* than profiles from 2D cultures[8]. A sharp increase in the number of publications on 3D cell cultures has been seen over recent years [9]. Pre-clinical *in vivo* animal models are often used in cancer research to test drug toxicity and new treatment regimens. The development of 3D models supports the reduction in the use of animal models as spheroids and organoids allow scientists to model tumours and organs for pre-clinical large scale treatment screens.

### Drawbacks of spheroids and organoids models

Despite the numerous advantages, spheroids and organoids so far are not widely used in biomedical research due to a lack of suitably high-throughput assays and reliable analysis methods [8]. Many spheroid assays are endpoint only, meaning that changes occurring between initial set up and the assay end point are not recorded. Evaluation of spheroid measurement changes over time including parameters such as area, diameter, and circularity could be a valuable tool for many biological studies. In many cases, obtaining these results involves manual imaging of each spheroid with subsequent manual application of imaging tools to obtain quantification metrics that are individually collated and then laboriously analysed. This is time-intensive, resulting in fewer technical repeats per condition being studied, and introduces subjectivity and, thus, human error. To address these issues, we have created SpheroidAnalyseR: an R Shiny app that takes spheroid size measurements and associated metadata as input, identifies and optionally removes outliers using a Robust-Z-Score, can merge datasets from multiple plates (including across a time-course) and can plot graphs to enable users to analyse spheroid data quickly and efficiently.

SpheroidAnalyseR identifies outliers in the data, including natural outliers and those caused by erroneous dimensional parameters for example those resulting from incorrect detection of the spheroid by the imaging software, or wells with missing spheroids due to human error. To maintain the quantitative integrity of the measurements it is therefore necessary to identify and remove any outliers from the dataset, which may compromise the validity of the final analysis. A configurable Robust-Z-Score used to measure outlier strength has been incorporated into SpheroidAnalyseR’s outlier detection code using statistical summarisation of the parameter data grouped by technical replicates [10].

### The R Shiny package

The R programming language is a highly regarded open-source platform for data manipulation, visualisation and statistical analysis. Despite numerous training initiatives, there is a still a steep learning curve for those without prior programming experience to adopt the language in their daily work. Shiny allows R developers to make their code and analyses accessible to the wider community through a web interface.

## PROCEDURE

The starting point for using SpheroidAnalyseR input of size metrics from spheroids/organoids grown in 96-well plates. This requires two prior steps:

1. Growing the spheroids in 96-well plates. This can be done via the user’s preferred method. Herein we used Ultra-low-attachment 96-well plates (Corning®, New York, US; catalogue number 7007).
2. Quantification of spheroid images.

There are multiple ways in which to acquire the input data for SpheroidAnalyseR. This includes manual imaging and subsequent measuring for example using an EVOS fluorescent and transmitted light microscope capable of producing high resolution images. ImageJ, a Java-based image processing programme, can be used to manually draw around each spheroid[11]. Next, data can then be uploaded into the template spreadsheet provided, and analysed by SpheroidAnalyseR. Alternatively, published methods such as SpheroidSizer provide a way to calculate the area and other key spheroid measurements from images [12]. In addition, we collaborated with Nikon to create a bespoke JOB (computationally scripted series of commands) for the Confocal Laser Scanning Microscope-Nikon A1R for use with spheroids grown in 96-well plates. This JOB performs spheroid imaging and subsequent measurement of volume, area, diameter, circularity and perimeter automatically using a threshold-based method, with results output to a Microsoft Excel file. A full 96-well plate is imaged, and result outputs are obtained within 2 minutes. The JOB can be accessed via the Glioma Genomic GitHub (https://github.com/GliomaGenomics) with full instructions on how to upload to and run on a Nikon A1R. The resulting Excel spreadsheet can be uploaded directly to SpheroidAnalyseR where users can preview the data, remove outliers, merge result files and plot graphs. The workflow for analysing the data is as follows:

### 1. Data input

SpheroidAnalyseR identifies and visualises outliers in spheroid data giving the user the option to remove some or all of them. It does this by converting user-defined technical replicate data into Robust-Z-Scores and allowing the user to set a threshold score, with the default set as ± 1.96, which equates to a 95% confidence interval. The webpage, after data input, includes graphical visualisation of spheroid measurements across multiple predefined parameters i.e., time, cell-type and treatment(s).

1.1 On the *Data Input* tab, three types of files are required to be uploaded to allow processing (see below). Input file templates that include the specific column headings needed can be downloaded from the *Data Input* tab.
  A. **Raw data file** - Files can be uploaded either directly from the output file from the bespoke JOB on the confocal (raw data template 1) or, data can be manually inputted into the second template file (raw data template 2). Several parameters can be inputted included treatment date and time, cell line, passage number, drugs and concentrations used, and irradiation dose. Any columns in which no data is present should have ‘0’ for each row. If the user is analysing multiple plates of data with the same layout, then multiple raw data files can be uploaded simultaneously to allow for faster processing.
  B. **Plate layout file** - This defines the layout of treatments on the 96-well plate for each spheroid or organoid. Each plate can contain spheroids or organoids with multiple different treatments. 1-12 represents columns 1-12 of the 96-well plate and A-H represents rows A-H of a 96-well plate. A number corresponding to the treatment index assigned in the treatment file should be in each cell that a spheroid measurement was for. Only one plate layout can be uploaded at any one time. If a raw data file has a different plate layout to the last file analysed, then a new plate layout will need to be uploaded.
  C. **Treatment file** - This file defines the treatment index numbers corresponding to the plate layout file. An Index denotes a specific treatment, or combination thereof, and cell line as detailed in the corresponding row of the treatment template. If desired, users can input the time and date of treatment, the cell lines used, the passage number, the dose of radiation used and multiple drugs and subsequent concentrations. Multiple wells with the same index constitute technical replicates. Multiple different combinations of treatments can be defined as required.

#### Note

Templates for these files are available to download on the *Data Input* tab of the SpheroidAnalyseR web page. The column names of any input files must match those given in the supplied templates, and the sheet name must match the template’s sheet name. Four template sheets are provided:

A. Raw data template 1 - this is the file created after running the bespoke JOB on the Confocal Nikon AR1. It can be uploaded straight to SpheroidAnalyseR.
B. Raw data template 2 – this template allows users to input their own spheroid measurements obtained. Users must have the Well.Name column completed and at least one of measurements from the other columns which are: area, perimeter, circularity, count (number of spheroids per well), diameter and volume. If any measurements are not inputted, then a ‘0’ should be inputted in each cell. **Optional**: – the user can input the time and date the spheroids were analysed, if not required then ‘0’ should be inputted in each cell.
C. Plate layout template.
D. Treatment template.

#### Caution

The number of spheroids to be analysed in the raw data file must match the number of cells filled out in the plate layout or an error will be shown in red on the *Outlier Removal* tab and the user will not be allowed to process outliers.

### 2. Previewing files in the *Data Input* tab

2.1 Raw data files can be previewed on the *Data Input* tab. A drop-down list enables users to select each raw file if multiple files were uploaded simultaneously. Treatment and layout information based on the templates uploaded can also be reviewed here: the plate layout is shown, and a dropdown list allows the user to colour the plate according to each parameter in the treatment file (i.e., cell line, drug used, or time treated).

#### Note

an image of the plate layout along with its designated treatment index is shown. It is recommended to review your uploaded data to ensure that that plate layout and treatment index are correct before proceeding.

### 3. Outlier removal

Once data has been inputted and reviewed, the user can move to the *Outlier Removal* tab and determine if any of the data points are spurious and should be removed as an outlier

3.1 On the *Outlier Removal* tab, choose the raw file to be analysed from the ‘Chose a raw file’ drop-down list.
3.2 3.2 A spheroid measurement value (e.g., diameter, area, etc.) for which the outliers should be determined for can be selected from the ‘Choose a value to plot the outliers’ drop-down list.
3.3 In the shaded beige/yellow box (Figure 2) users can adjust the Robust-Z-Score (if required) and choose to apply pre-screen thresholds. These are automatically applied with default settings but can be adjusted if necessary. The pre-screen threshold removes spheroid measurements where the value falls outside the upper or lower limits set by the user. This ensures any wells in which there is an extreme outlier, for example due to an empty well will be removed prior to outlier analysis.
3.4 After clicking the ‘Remove outliers’ button, outliers will be removed, and three images will be displayed. The top figure shows the plate layout after wells with spheroids that are determined to be outliers have been removed based on the pre-screen thresholds and Robust-Z-Score limits. Below that, Plot 1 and Plot 2 can be viewed with the selected measurement (e.g., diameter, area, etc.). These plots show the columns of the plate and the measurement each spheroid in these columns has (for example diameter). Plot 2 shows spheroids grouped by technical replicates, i.e., with the same treatment index as defined on the previously inputted treatment file.

#### Hint

It is advised to apply pre-screen threshold to remove any values that are obvious outliers before running the Robust-Z-Score. This could include measurements that have been taken of empty wells or when the imaging technique failed to recognise and measure a spheroid.

3.5 Using the ‘Toggle Cell Status normal/outliers’ drop-down list, users can manually override individual results of the outlier process. The user must select a cell or multiple cells and choose the ‘apply manual adjustment’ button. At this point, if the spheroid in that well was determined to be an outlier, it will now be classed as a normal result, and vice-versa. This is to give the user full control over the data and inclusion in subsequent analysis.
3.6 The report of the selected file with outliers removed can be downloaded by clicking the Download button. This will automatically download and be titled <the file name of the raw file>. The file contains multiple tabs each showing a different measurement (area, diameter etc.) with two plots per tab showing the data for each treatment index with and without outliers. The main dataset tab shows all the data uploaded with new columns added with the outliers removed (OR) (Area_OR etc). This data can be used by the user to create their own graphs and perform statistical analysis in their preferred method. The downloaded files can also be re-uploaded later to the *Merging* tab by selecting ‘Use previous reports’ checkbox and using the file selection tool to upload the previously processed files.

### 4. Merging

4.1 The *Merging* tab allows multiple files to be merged into one master file to create plots in the *Plotting* tab. If the user is only analysing one file, then the ‘Merge’ button should be chosen and then progress straight to the next tab.
4.2 On the main panel on the *Merging* tab, a table of raw files that have been uploaded on the *Data Input* tab will be shown. The third column (column name: Processed) shows whether the file has been processed through the outlier removal step on the *Outlier Removal* tab. The remaining columns show the configuration (e.g., Robust-Z-Score value or Pre-screen thresholds) used in outlier removal so that the user has a record of the parameters should they need to re-analyse a file or analyse future files in the same way later.
4.3 Files that have been previously analysed on the *Outlier Removal* tab can be uploaded directly to the *Merging* tab. To do this, select the ‘Use previous reports’ checkbox and browse for the correct files.
4.4 The merged file is only available when selected raw files have been processed. Once all the raw files have been processed on the *Outlier Removal* tab, press the Merge button. This will allow plots to be created using the data from multiple files on the *Plotting* tab. The user can rename (using the ‘Merged file name’ textbox) and download (using ‘Download the merged file’ button) the merged file if required.

### 5. Plotting

5.1 Plots of the merged data can be created and viewed on the *Plotting* tab. SpheroidAnalyseR supports bar charts, point plots, dot plots, and boxplots. The user must select a plot type (bar chart by default) followed by a spheroid/organoid measurement type for the Y-axis (Area by default) and a grouping parameter for the X-axis (Treatment.Label by default) from the drop-down lists. Up to two grouping parameters can be specified to be distinguished by different colouring.
5.2 The plot can be named and there is an option to colour the plot in black and white. Press the ‘Plot’ button to view the plot on the tab, and the ‘Add to the report’ button to add it to the report which can be downloaded using the ‘Download the report of plots’ button. Multiple plots can be added to the report before downloading.

## ANTICIPATED RESULTS

### Example work through

A series of screenshots taken from SpheroidAnalyseR highlighting the process of uploading data, data analysis and data presentation is given in Figures 1–4. Key steps are described in the associated figure legends. The example data files are provided in the supplementary materials (raw data is in Supplementaryfile1, plate layout is supplied in Supplementaryfile2 and treatment definitions within Supplementaryfile3).

**Figure 1.**
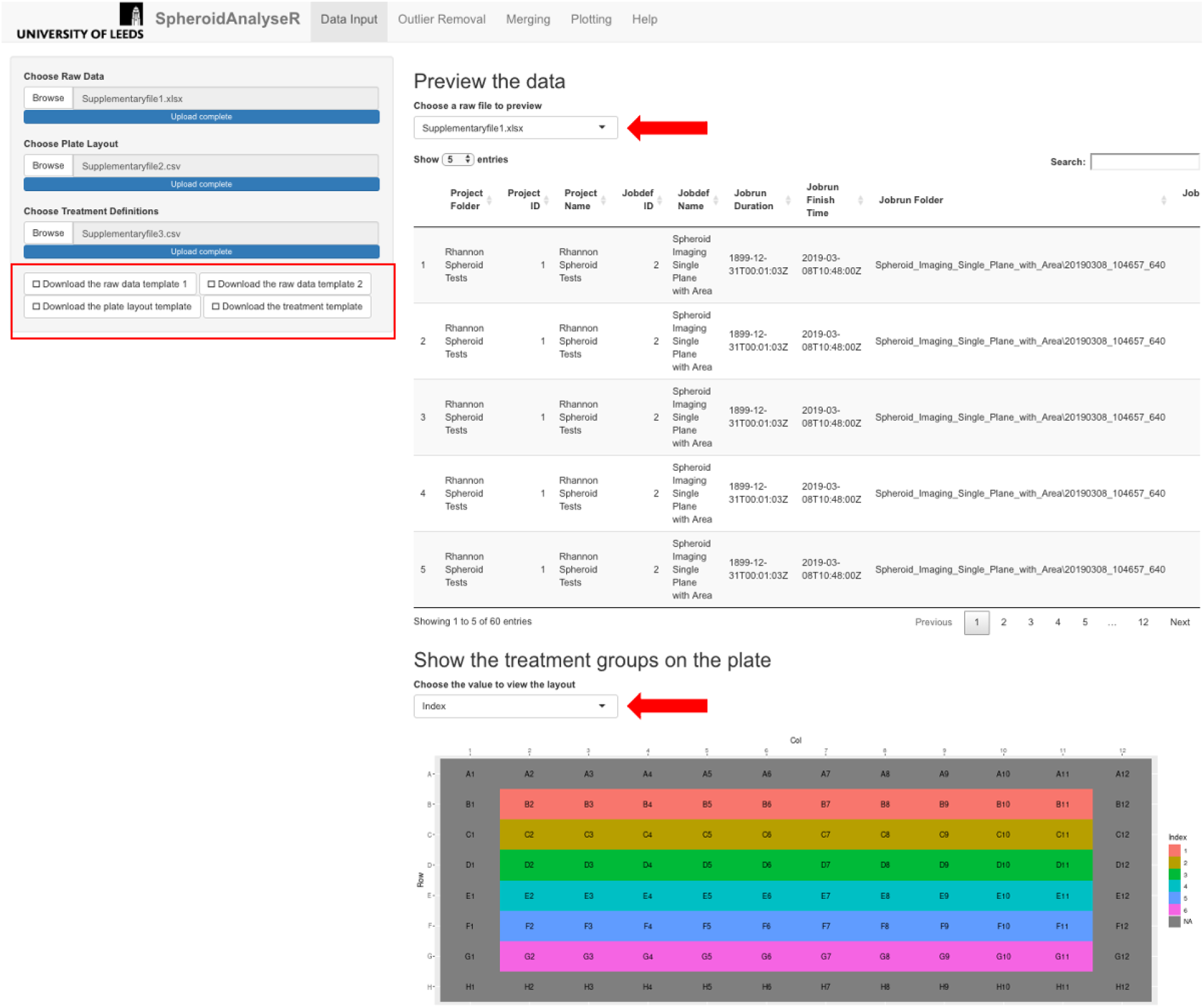
The *Data Input* tab once raw data has been uploaded. Template files are available to download (see red rectangle). A raw file (Supplementaryfile1), plate layout (Supplementaryfile2) and treatment file (Supplementaryfile3) have been uploaded. A preview of the raw data is shown in the main panel, and a map of the plate coloured by treatment index is displayed underneath. Red arrows show the dropdown menus to select a different file to view, and to choose a value to view the layout for.

**Figure 2.**
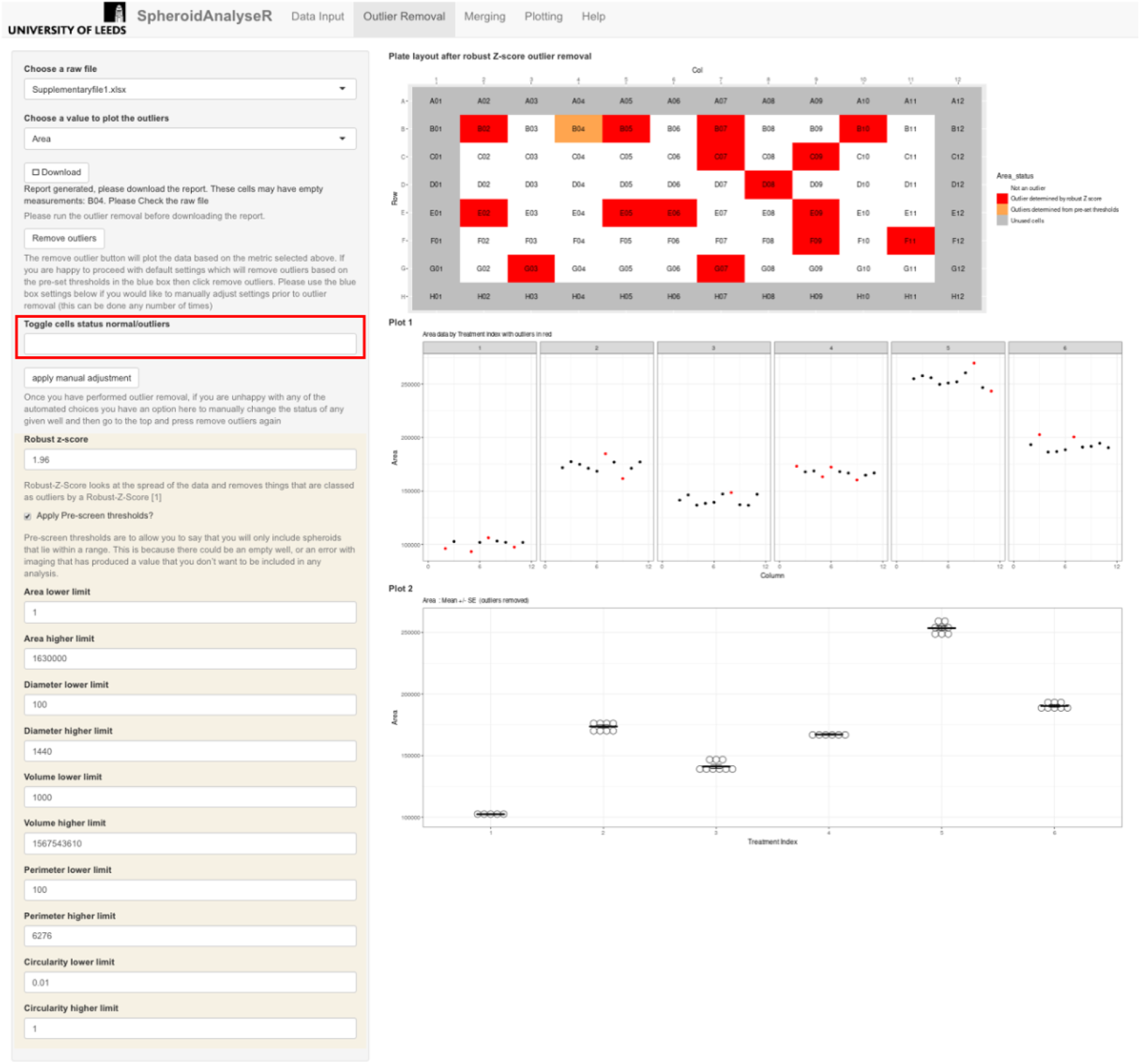
The *Outlier Removal* tab. Inside the beige box are the adjustable outlier settings of Pre-screen-thresholds and Robust-Z-Score. After pressing the ‘Remove outliers’ button, the outliers have been identified on the plate layout seen at the top of the panel. Orange shows outliers removed via pre-screen thresholds and red shows outliers removed after Robust-Z-Score. The results of either of these can be modified using the ‘Toggle cell status, normal/outlier’ drop down menu. The user should select the wells to adjust and press ‘Apply manual adjustment’. The data can be downloaded as an excel spreadsheet via the Download button.

**Figure 3.**
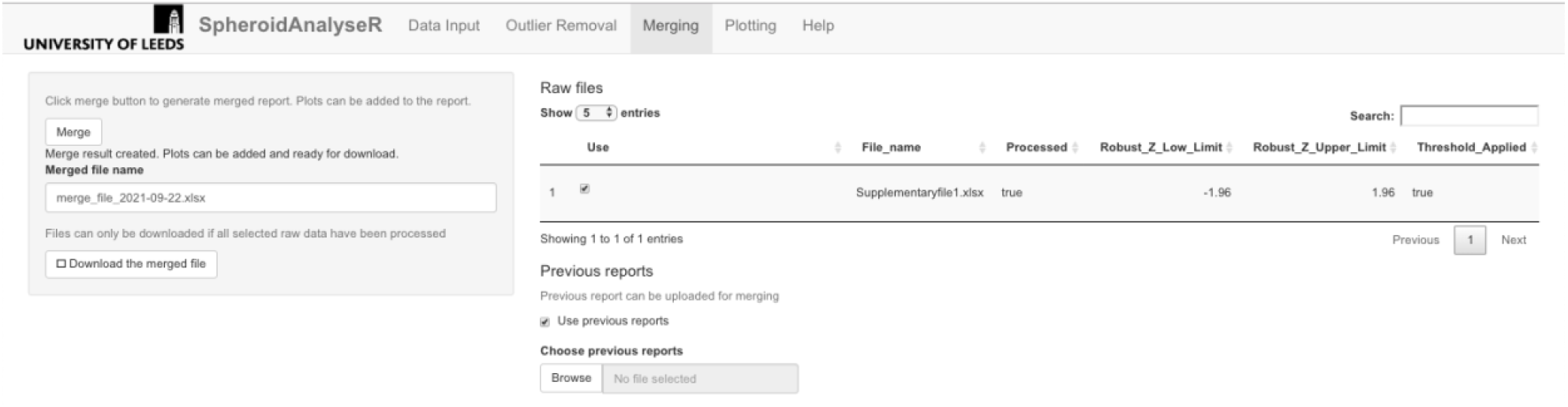
The *Merging* tab. The status of the file processing is shown ‘true’ or ‘false’ in the Processed column. The Merge button must be pressed before moving to the next stage even if there is only one file to be analysed. Files that have been downloaded from the ‘*Outlier Removal*’tab can be uploaded here if required.

**Figure 4.**
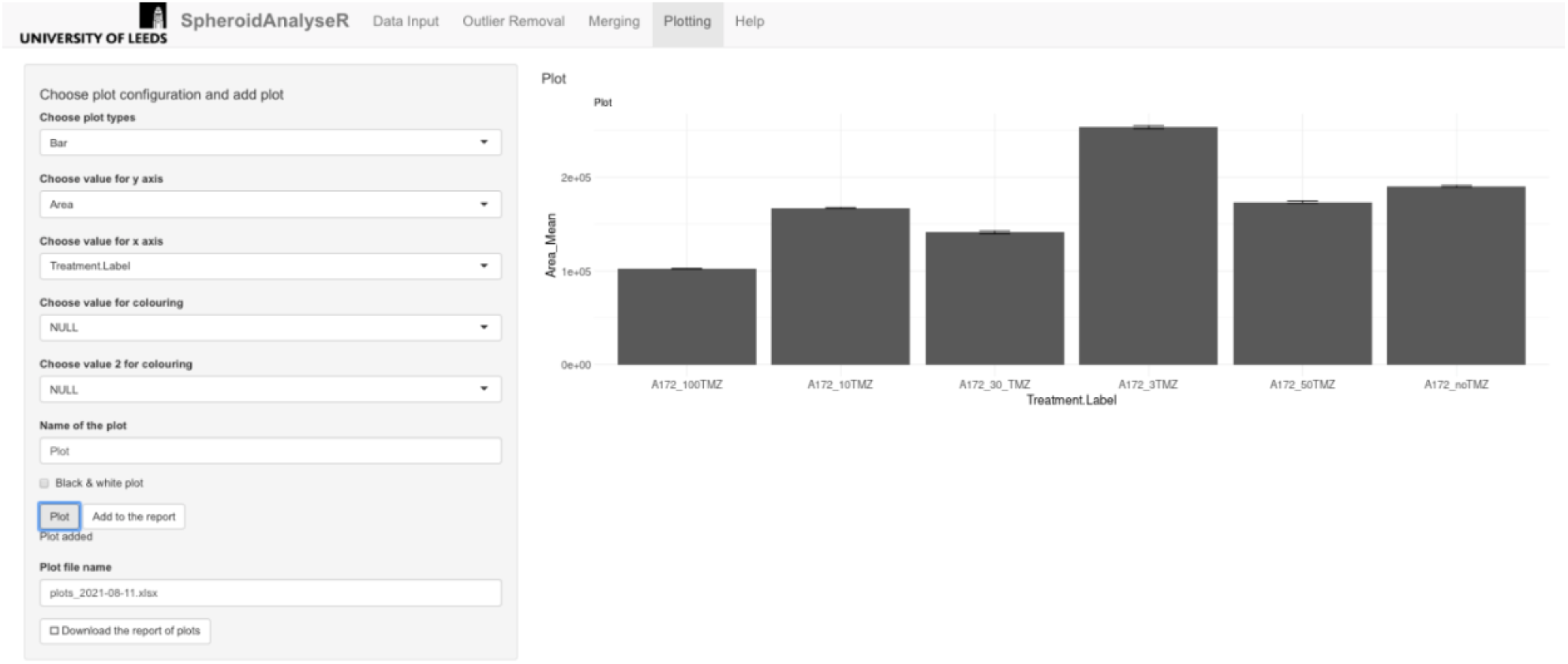
Shows the *Plotting* tab. The default plot type is a bar plot, but different plot types can be selected in the ‘Choose plot type’ drop-down list. Values to plot on the Y-axis and X-axis can be selected from the drop-down lists. The plot can be displayed in black and white if desired. Plots can be added to the report via the ‘Add to report’ button and then a file of the plots can be downloaded.

## TROUBLEHSOOTING

**Table 1.** Trouble shooting for the SpheroidAnalyseR app.

| Step | Problem | Cause | Solution |
| --- | --- | --- | --- |
| Outlier removal.<br>3.4 | Error message -<br>*Number of rows of data in raw file does not* | The number of rows of data in the raw file is | Check that the layout template has the correct |
|  | *match the plate setup checksum* | different to that specified in the layout template | number of spheroids labelled |
| Merging Tab.4.4 | Error message - *Please check all selected files have been processed* | Not all files uploaded to the app have been processed on the *Outlier Removal* tab | Using the select box on the *Outlier Removal* tab, go through each raw file available and press the remove outlier button for each file. |

## Supporting information

Data file example - from the confocal

Plate layout example

Treatment file example

## Notes

https://github.com/GliomaGenomics

https://spheroidanalyser7.azurewebsites.net

### Competing Interest Statement

The authors have declared no competing interest.

